# A food color based colorimetric assay for *Cryptococcus neoformans* laccase activity

**DOI:** 10.1101/2024.01.01.573823

**Authors:** Lia Sanchez Ramirez, Quigly Dragotakes, Arturo Casadevall

**Author notes:** Address correspondence to Lia Sanchez Ramirez or Arturo Casadevall, or.

## Abstract

*Cryptococcus neoformans* is a fungal pathogen that causes cryptococcosis mostly in immune compromised patients, such as those with HIV/AIDS. One survival mechanism of *C. neoformans* during infection is melanin production, which catalyzed by laccase, and protects fungal cells against immune attack. Hence comparative assessment of laccase activity is useful for characterizing cryptococcal strains. We serendipitously observed that culturing *C. neoformans* with food coloring resulted in the degradation of some dyes with phenolic structures. Consequently, we investigated the color changes for the food dyes metabolized by *C. neoformans* laccase and explored using this effect for the development of a colorimetric assay to measure laccase activity. We developed several versions of a food dye based colorimetric laccase assay that can be used to compare the relative laccase activities between different *C. neoformans* strains. We found that phenolic color degradation was glucose dependent, which may reflect changes in the reduction properties of the media. Our food color based colorimetric assay has several advantages over the commonly used 2,2′-azino-bis (3-ethylbenzothiazoline-6-sulfonic acid) (ABTS) assay for laccase activity, including lower cost, irreversibility, and does not require constant monitoring. This method has potential applications to bioremediation of water pollutants in addition to its use in determining laccase virulence factor expression.

**Importance:** *Cryptococcus neoformans* is present in the environment and while infection is common, disease occurs mostly in immunocompromised individuals. *C. neoformans* infection in the lungs results in symptoms like pneumonia, and consequently cryptococcal meningitis occurs if the fungal infection spreads to the brain. The laccase enzyme catalyzes the melanization reaction that serves as a virulence factor of *C. neoformans*. Developing a simple and less costly assay to determine laccase activity in *C. neoformans* strains can be useful for a variety of procedures ranging from studying the relative virulence of cryptococci to environmental pollution studies.

## Introduction

*C. neoformans* causes cryptococcosis and is a public health concern due to its high mortality rate in immunocompromised individuals, especially those with HIV/AIDS. Consequently, *C. neoformans* was recently designated as a critical priority pathogen by the World Health Organization. Most cases of cryptococcosis occur in Sub-Saharan Africa, where cryptococcal meningitis has become a major cause of death in HIV/AIDS patients, surpassing the mortality rate of tuberculosis^1^. Therefore, *C. neoformans* infections are particularly dangerous in low-resource countries where these is low access to treatment for immunocompromised individuals. Studying the ways in which *C. neoformans* protects itself against an immune response allows us to further understand the immune defenses that combat *C. neoformans* infection.

Laccases catalyze a broad spectrum of reactions, and the enzyme serves as an important virulence factor for *C. neoformans* that is expressed in its cell wall^2^. Since laccases are versatile in the sense that they can catalyze the oxidation of various substrates, this makes them useful for a range of environmental applications. Laccases can oxidize polyphenolic compounds and iron, and the functions of the enzyme are regulated through signal transduction pathways^2^. Laccase has even more diverse functions, since a broad range of organisms utilize the laccase enzyme including fungi, plants, and insects^2^.

Specifically in *C. neoformans*, the laccase enzyme convers Iron (II) to Iron (III) which lowers the susceptibility of *C. neoformans* cells to hydroxyl radicals that are generated in vitro^3^. Regulation of laccase expression has been found to possibly be strain-dependent, because disruptions of a G-protein subunit homolog affect laccase activity in wild-type *C. neoformans* but this was not observed in a serotype A H99 strain^2^. More research is needed to understand the molecular complexities of the laccase enzyme serving as a virulence factor of *C. neoformans*. Vacuolar proton pump ATPases were found to play a role in laccase activity, by testing strains with mutated genes that encode for ATPases, so targeting these proton pumps could be useful for drugs that aim to control laccase activity^2^.

Laccase fills an important role in virulence by catalyzing the generation of melanin in *C. neoformans*, which protects the fungus from reactive oxygen and nitrogen oxidants produced by phagocytic^3^. Fungal laccase may also contribute to virulence by reducing the formation of antimicrobial hydroxyl radicals^3^. Laccase is also involved in the generation of prostaglandins by the fungal cell that can affect local inflammatory responses^4^. Specifically, laccase is induced during glucose starvation and increased temperature (30 °C) and helps moderate fungal stress resistance^2^. Studying laccase can provide insight into *C. neoformans* pathogenesis and its mechanisms of defense in the phagolysosome. The *C. neoformans* laccase is known to have broad structural activity producing pigments of a spectrum of colors, including those similar to melanin, following the oxidation of phenols and cetechols^5^.

Laccases are known to destroy dyes and have uses in food preparation, industry, and environmental applications such as reducing the environmental impact of synthetic dyes^6, 7^. Factory production of non-biodegradable synthetic dyes results in contaminated wastewater and fungal laccases can be used to degrade the colors in these bodies of water^6^. The by-products of laccase catalysis are lower in toxicity than alternative chemical methods for wastewater purification and laccases have broad specificity, which allows them to break down a variety of synthetic dyes^6^. Additionally, a common substrate for laccase is phenol, a toxic pollutant which is dangerous to humans^6^. The use of laccases as versatile agents of bioremediation is emerging as a sustainable option to degrade chemical pollutants, and there is interest in investigating the catalytic mechanisms of laccase to optimize its function in bioremediation^8^.

In this study, we followed up a serendipitous observation to investigate the use of mangosteen-colored food dye in a fast, efficient, low-cost colorimetric assay of laccase activity in *C. neoformans* cultures. Using absorbance spectroscopy, we were able to compare several strains of *C. neoformans* in various culture conditions to both detect and quantify relative laccase activity. Low cost and facile assays for laccase activity have many potential applications in environmental industries, including purification of wastewater. These assays are also applicable when comparing the virulence of different strains by measuring relative laccase activity, an important fungal virulence factor.

## Methods

### Yeast Strains and Culture Conditions

*C. neoformans* species complex serotype A strain H99 was obtained from John Perfect in Durham, NC.^9^ The *lac1*Δ mutant is from the 2007 Lodge library (Fungal Genetics Stock Center)^10^. The H99 GFP strain was obtained from the lab of Dr. Robin May at the University of Birmingham, United Kingdom^11^. The CNAG 01373Δ, CNAG 06646Δ, and CNAG 01029Δ strains are KN99α mutants obtained from a previously published gene knockout library^12^. KN99α strain obtained from Heitman Lab at Duke University Medical Center Durham, North Carolina^13^. Additional *C. neoformans* strains used are 24067, Mu-1, and B3501. Other yeast strains used for comparisons were *C. albicans* 90067, *S. cerevisiae* S188C, *C. gatti* R265, and *C. gatti* WM179.

### Media preparation

Minimal media (MM) was prepared as 15 mM dextrose, 10 mM MgSO_4_, 29.3 mM KH_2_PO_4_, 13 mM glycine, and 3 μM thymine-HCL dissolved in water supplemented with either 2.7 g/L or 20 g/L glucose, then vacuum sterilized via SteriCup Quick Release filter and stored at room temperature. Yeast Extract Peptone Dextrose (YPD) was prepared according to manufacturer protocol.

### Comparison of color change in H99 and lac1Δ mutant cultures

H99 and *lac1*Δ mutant strains were seeded in 3 mL of MM in 12-well tissue culture plates. The following colors of Limino brand food coloring (Limino Baiyun, Guangzhou, China) were used: Strawberry, Tangerine, Lemon, Lime, Purple Cabbage, Blueberry, and Mangosteen. Each well had a concentration of 10^4^ cells /mL, with 10 wells containing MM supplemented with 10 µL food coloring, one well with uncolored MM, and one well with uncolored YPD. Color change observations were recorded 7 d after the plate was either left on the bench at room temperature or placed on a 120-rpm shaker in a 30°C incubator and measured either by eye or via Spectramax iD5.

### Different concentrations of glucose were tested with high glucose minimal media conditions

Two H99 cultures were seeded in MM supplemented with mangosteen color in 17 mL polystyrene culture tubes. One tube was prepared with MM at its regular glucose concentration of 2.7 g/L, while the second tube was prepared with MM at an elevated glucose concentration of 20 g/L. Culture tubes were placed in the 30°C incubator with rotation and color change observations were recorded after 7 d.

### Kinetic Assay for Laccase Activity

24-well tissue culture plates were seeded with 10^6^ *C. neoformans* cells in 1 mL of MM with 10 g / L glucose in each well. Three wells were immediately treated with Thermofisher antioxidant reagent (Product Number: NP0005), then the plate was incubated at 30°C shaking overnight. The next morning, antioxidant reagent was added to the last three wells and incubated for another hour to observe possible color change of blue to green or revert to purple.

### Addition of antioxidant to colorimetric assay

96-well tissue culture plates were seeded with 10^4^ *C. neoformans* cells in 100 μL volumes of MM. The 1:1000 working dilution of antioxidant was added either at 0 h or after 24 h of observing the color change.

### 24-hour Assay for Laccase Activity

Every culture of interest was grown in regular MM and left in a 30°C incubator with rotation. A 1.5 mL volume of each culture was centrifuged at 2300 g for 5 minutes in microcentrifuge tubes, and then resuspended in 1.5 mL of 10 g/L MM. 150 μL of the resuspended culture was pipetted into 10 wells of a 96-well plate, so each culture was measured with 10 replicates of the well. Then, 7.5 μL of a 1 to 10 dilution of Mangosteen food coloring in water (100 μL food coloring in 1 mL of H_2_O) was added to each well. At 0 h, 50 μL of supernatant were placed in another 96- well plate for the initial 520 nm absorbance measurement with the Spectramax iD5 (Baltimore, Maryland) and at 24 h another 50 μL of supernatant were placed in another 96-well plate for the 24h measurement with the Spectramax iD5. For those 24 h, the plate was incubated in a 30°C incubator on 120-rpm shaker. We compared the laccase activity of the following *C. neoformans* strains: H99, H99 GFP, KN99α, CNAG 01373Δ, CNAG 06646Δ, and CNAG 01029Δ. Additional trials were conducted comparing the following *C. neoformans* strains: H99, 24067, Mu-1, and B3501 to the following other yeast strains: *C. albicans* 90067, *S. cerevisiae* S188C, *C. gatti* R265, and *C. gatti* WM179.

### 2,2′-azino-bis (3-ethylbenzothiazoline-6-sulfonic acid) (ABTS) assay

H99 strain was seeded in MM cultures. Each culture used was washed twice with phosphate buffered saline (PBS). Then a 1:100 dilution, or 1:1 dilution for some wells, was prepared with MM. A 1 mL volume of 20 mM of ABTS solution was prepared and filter sterilized. ABTS was added for final concentration of 1 mM ABTS in the cultures. Incubate for 24 h in 30°C incubator on 120rpm shaker. Initial and 24 h absorbance measurements were taken at 734 nm with the Spectramax iD5.

### Statistical Analysis

The statistical tests conducted for each absorbance measurement experiment are denoted in their respective figure descriptions with tests for multiple hypotheses. Two-way ANOVA with Tukey comparison analyses were conducted using RStudio Version 2023.09.1+494 and GraphPad Prism Version 10.0.2(171). Statistical comparisons were made both with all the data for each individual experiment replicate and for data from all trials pooled.

## Results

### Phenolic Dyes are Degraded in C. neoformans Culture

The observation that *C. neoformans* degrade some food dyes was made serendipitously. While searching for conditions to study the growth of *C. neoformans,* we noted difficulty in the measurement of fungal growth by turbidity when comparing cultures grown in MM and YPD. We hypothesized that the problem arose because MM was clear while YPD had a yellowish color and thought that adding food coloring to MM wells would facilitate the interpretation of growth curve absorbance data. However, after 7 d of culture growth, we noticed a color change occurred in multiple wells. Upon closer inspection, we noted color changes only in dyes containing a red component, whose chemical structures contained phenolic groups, and that the resulting colors resembled the original dye if only the phenolic (red) component were removed **(Figures 1 and 2A)**. When absorbance was measured, we observed a loss of the red color in wells with *C. neoformans* culture that had degraded the red dye (**Figure 2E)**. Specifically, these wells contained the phenolic dyes Food Red 7 and Acid Red 27 **(Supplemental Table I)**. We hypothesized that the dye degradation was a result of laccase activity, due to broad substrate specificity in cryptococcal laccase, and established this by replicating the experiment with Δ*Lac1*-H99, and found no color change after 7 d^14, 15^. Taken together, these data suggest that fungal laccase can degrade phenolic food dye and we hypothesized this phenomenon could be harnessed for a colorimetric laccase activity assay.

**Figure 1.**
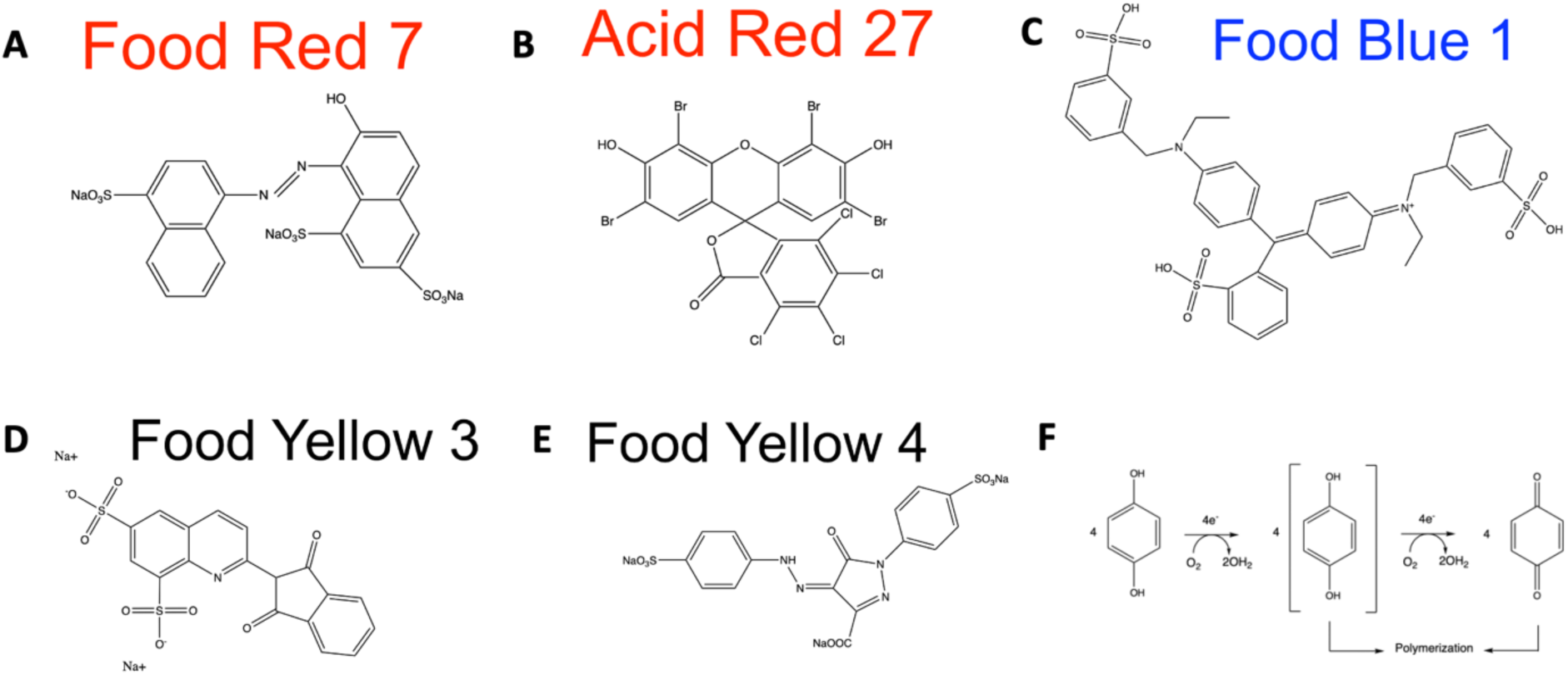
Chemical structures of food dyes used and melanization mechanism Panels A and E drawn from World dye variety^29, 30^ with ChemDraw software (Version 20.1.0.112)^31^, Panels B, C, and D adapted from NIH PubChem^32–34^, Panel F from Chandrakant and Shwetha^35^ and redrawn using ChemDraw software (Version 20.1.0.112)^31^. **A.** Food Red 7 **B.** Acid Red 27 **C.** Food Blue 1 **D.** Food Yellow 3 **E.** Food Yellow 4 **F.** Oxidation by laccase mechanism

**Figure 2.**
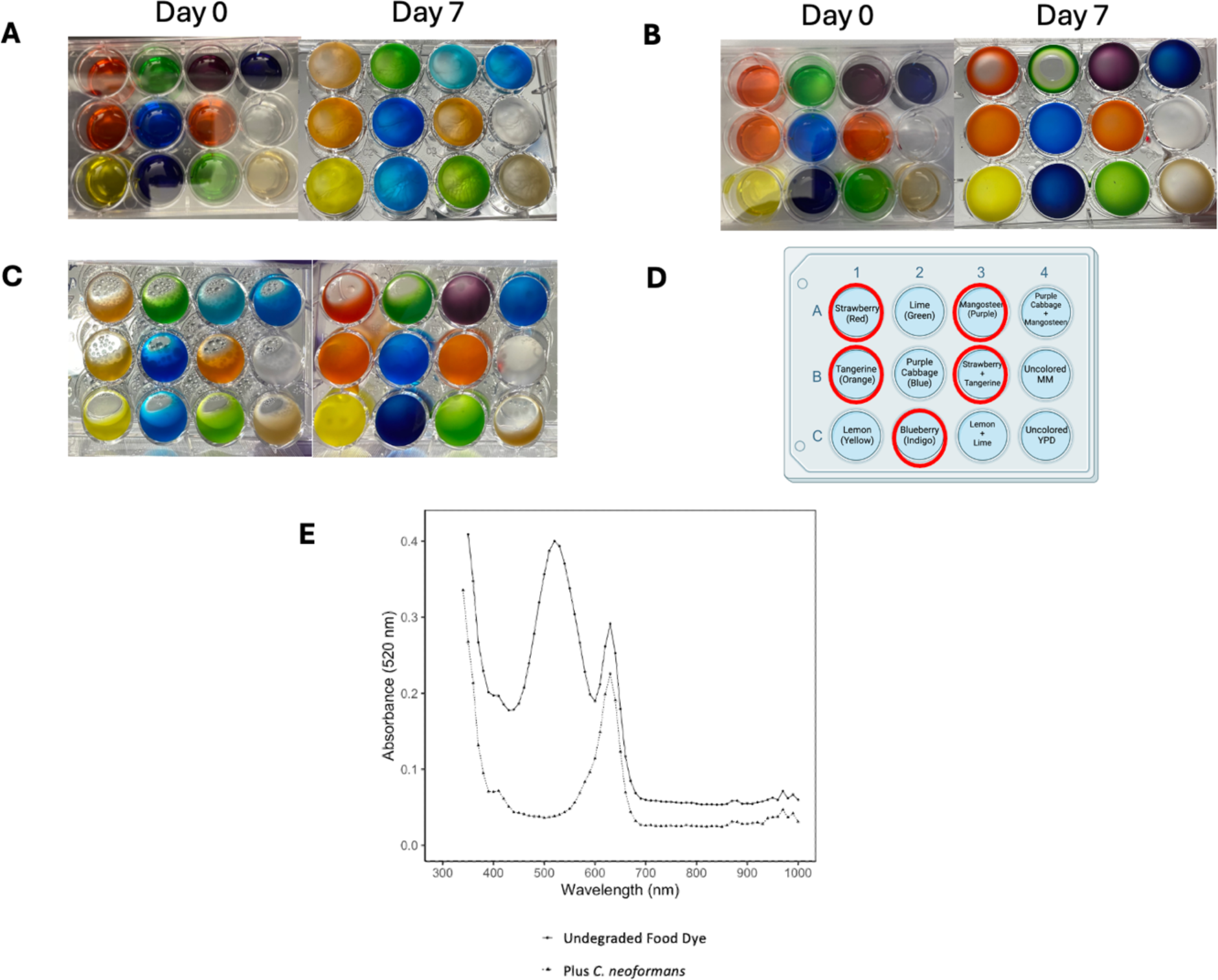
Observed degradation of red color after 7 days in different temperature, agitation, and culture conditions. **A.** Left: Plate with wild-type H99 culture. Right: After 7 days of agitation at room temperature. **B.** Left: Plate with *lac1*Δ mutant culture. Right: After 7 days of agitation at 30°C. **C.** Left: Wild-type H99 plate after 7 days of agitation at 30°C. Right: Wild-type H99 plate after 7 days left at the bench. **D.** Sample 12-well plate setup showing the location of the various compounds in the assay wells shown in panels A to C. Wells with a red circle designation are colors that commonly showed the red color degradation. Image created with Biorender.com. **E.** Absorbance measurements showing loss of red color peak in wells with *C. neoformans* culture compared to wells with undegraded food dye. Experiments were repeated multiple times with similar results.

### Glucose Concentration, Culture Agitation, and Cell Density Affect Laccase Activity

We sought to optimize culture conditions for detecting laccase expression and activity, attempting to determine conditions which would provide robust results while remaining cost and time efficient for any basic laboratory. We found that a 7 d incubation with agitation at 30°C was required to observe color change with a standard MM preparation **(Figure 2C)**.

We observed that high glucose concentrations (10-20 g/L) induced color change even at room temperature and without agitation at high initial cell densities. Additionally, using glucose concentrations of 10 g/L and 20 g/L MM allowed us to observe a visible color change quicker than regular glucose MM with any cell concentration from 10^3^-10^7^ cells / mL in a plate with agitation at 30 °C **(Figure 3A).** We found that high glucose concentrations repressed laccase expression **(Figure 3B).** Through an examination of glucose-dependence, we observed increased dye degradation in wells with cultures resuspended in MM with higher glucose concentrations **(Figure 4A).** Using a higher glucose concentration allowed us to view color change effects quicker as compared to wells with lower glucose concentrations, and with a 24 h assay we were not able to see any color change occur in MM glucose condition **(Figure 4B).**

**Figure 3.**
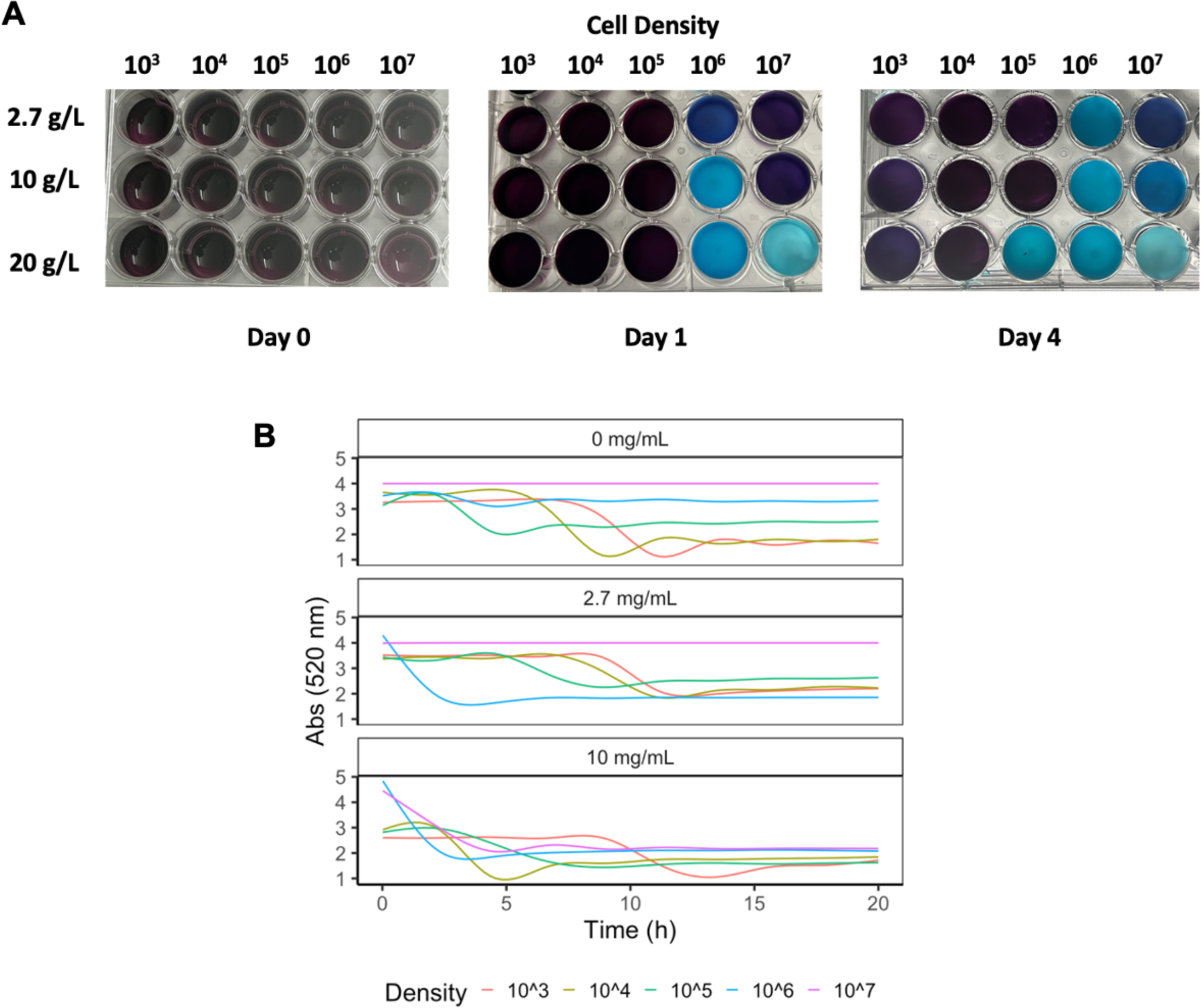
**A.** Mangosteen (Purple) to blue color change observed in wells with varying cell densities with regular 2.7 g/L MM, MM with 10 g/L of glucose, and MM with 20 g/L of glucose. Experiment conducted for two trials. **B.** Absorbance measurements at 520 nm wavelength for wells with varying cell densities and MM glucose concentrations.

**Figure 4.**
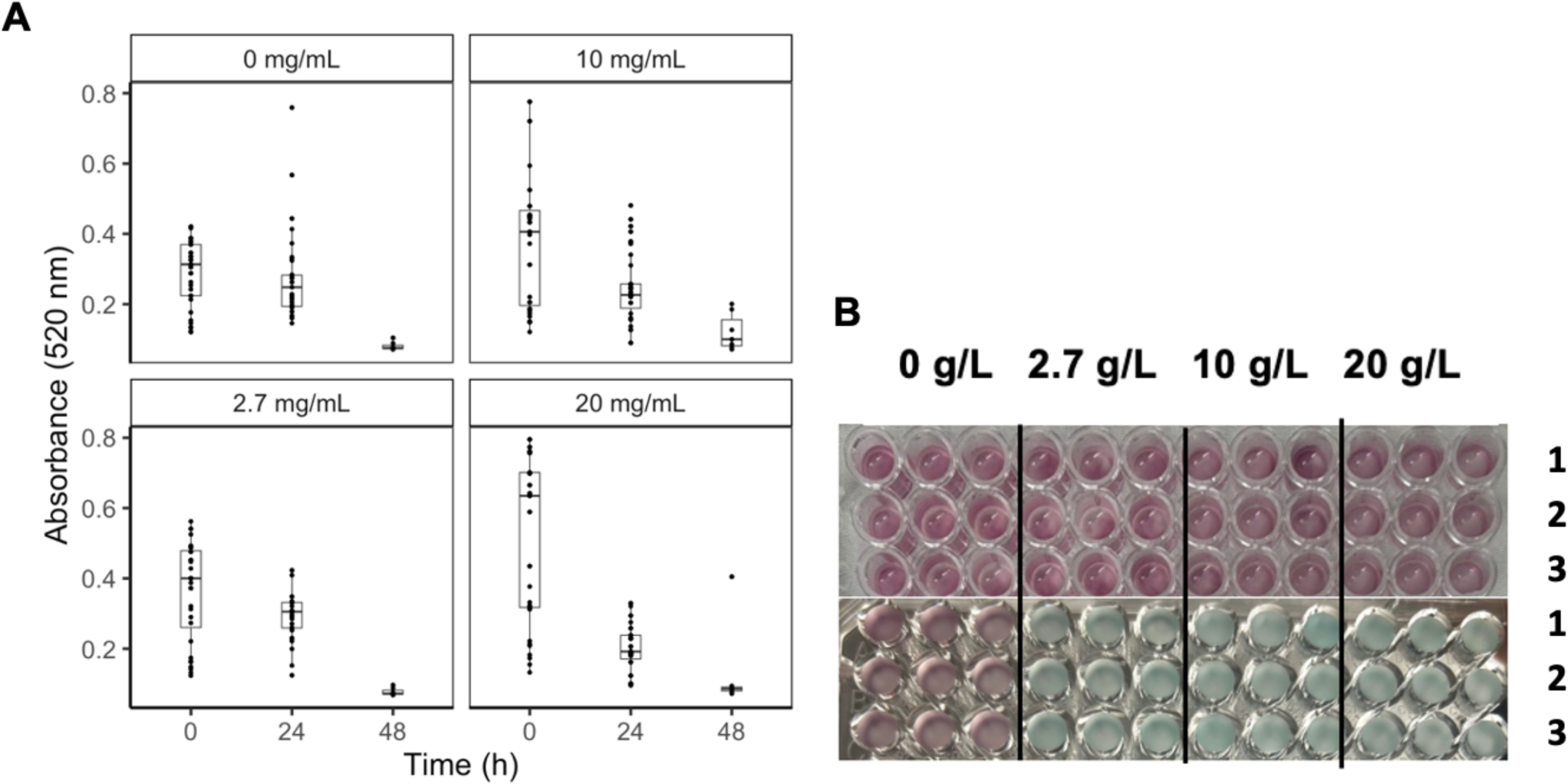
**A.** Absorbance measurements of supernatant at 520 nm at 0 and 24 h for different glucose concentrations. Each point represents an individual measurement. The following significant differences were determined when analyzing difference in absorbance across different glucose conditions over 24 h: 10 g/L and 0 g/L condition (P = 0.0002), 20 g/L and 0 g/L condition (P << 0.00001), 20 g/L and 10 g/L condition (P<<0.00001), and 20 g/L and 2.7 g/L condition ((P<<0.00001). Experiments conducted in triplicate. **B.** Mangosteen (Purple) to blue color change observations. No color change observed with 0 g/L glucose MM in the 24 h colorimetric assay. Rows 1-3 are replicates of the given glucose condition with the H99 strain. Each experiment is conducted in triplicate, vertical columns

With the combined data containing all individual trials of the glucose dependence experiments measuring absorbance at 520 nm, a statistically significant difference was found between the 20 g/L glucose MM condition when compared to the 0 g/L glucose MM condition (P = 0.002). An ANOVA test for glucose dependence was also conducted with the change in absorbance measurements between 24 h and 0 h of exposure to food coloring, which revealed significant differences between the 10 g/L and 0 g/L conditions (P = 0.0002), 20 g/L and 0 g/L conditions (P << 0.00001), 20 g/L and 10 g/L conditions (P<<0.00001), and 20 g/L and 2.7 g/L conditions ((P<<0.00001). Although not many significant differences were observed when analyzing the absorbance measurements themselves, analyzing the change in absorbance throughout the 24- hour timeframe showed differences between the glucose conditions. Overall, it was easier qualitatively to observe differences in laccase activity across different glucose conditions than it was to establish the quantitative differences which we suspect could be due to the sensitivity of equipment used.

### Laccase Dye Degradation is Irreversible

One potential disadvantage of current laccase activity assays, and specifically with the commonly used ABTS assay, is that the observed color change is not permanent and must be measured within a specific timeframe. To investigate whether our colorimetric assay was permanent, we treated wells with commercially available Thermofisher antioxidant before and after the observed color change. We found that adding antioxidant after the reaction did not revert the color, nor did leaving the sample on a benchtop long term over a span of 3 weeks **(Figure 5A)**. This suggests that the reaction was irreversible, allowing samples to be read at the investigator’s convenience. Interestingly, when treating the culture with antioxidant at the start of the experiment, we observed a new green color. A spectrum scan revealed a new peak at ∼420 nm, suggesting that an alternative product was formed **(Figure 5B).**

**Figure 5.**
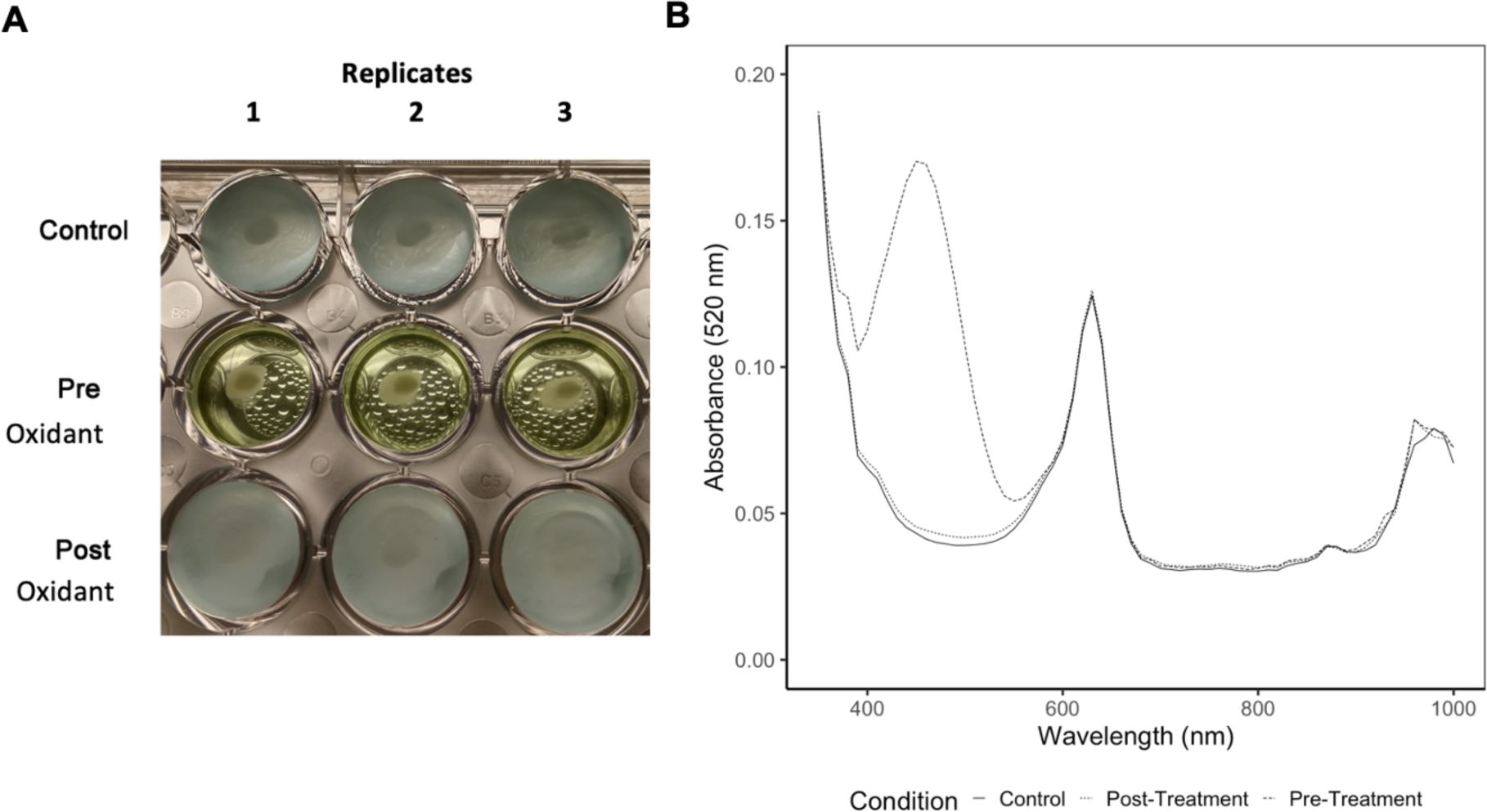
**A.** Thermofisher antioxidant treated wells. Top row is the control row without addition of antioxidant, middle row is addition of antioxidant before color change occurs, and the bottom row is addition of antioxidant after color change occurs. Each column is a replicate of the given condition. All wells were initially mangosteen (purple) color. The pre-oxidant row was given antioxidant before any color change occurred while the control and post oxidant rows had antioxidant added after they turned blue due to red color degradation. Color change is not reversible, since the wells turned green and not back to purple. **B.** Absorbance measurements at different wavelengths for wells that either degraded the food coloring to produce color change or did not degrade the coloring. Note that a new peak at ∼420 nm was observed in the absorbance measurements of wells with color degradation.

### Developing Cheap, Available, and Lightweight Laccase Activity Assay

Our next aim was to minimize the necessary reagents and instruments in this assay **(Supplemental Figure 1)**. The optimized assay protocol uses a concentration of 10^6^ cells / mL yeast seeded in MM with 7 uL of 1:100 diluted food coloring in a 96-well plate. The plate is then incubated at 30°C with 120-rpm agitation to observe color change in around 3-7 days.

To quantify the extent of red food coloring degradation in 24 h, we designed an assay using culture resuspended in 10 g/L MM. This quicker assay involves centrifuging the culture of interest and resuspending cells in 10 g/L glucose MM, with 100 μL of culture in each well in a 96-well plate and utilizing 7.5 μL of a 1 to 10 dilution of food coloring in water for each well. Absorbance measurements at 520 nm at 0 and 24 h can be used to compare differences between strains or other conditions to quantify changes in laccase activity **(Figure 6).** Linear regression analysis allowed us to compare rates of dye degradation across various *C. neoformans* strains, with increased rates of degradation in an H99 GFP strain (**Figure 6B and Figure 7**). Differences between multiple *C. neoformans* strains and other yeast strains with varying levels of laccase activity were compared with the 24 h assay **(Figure 8)**.

**Figure 6.**
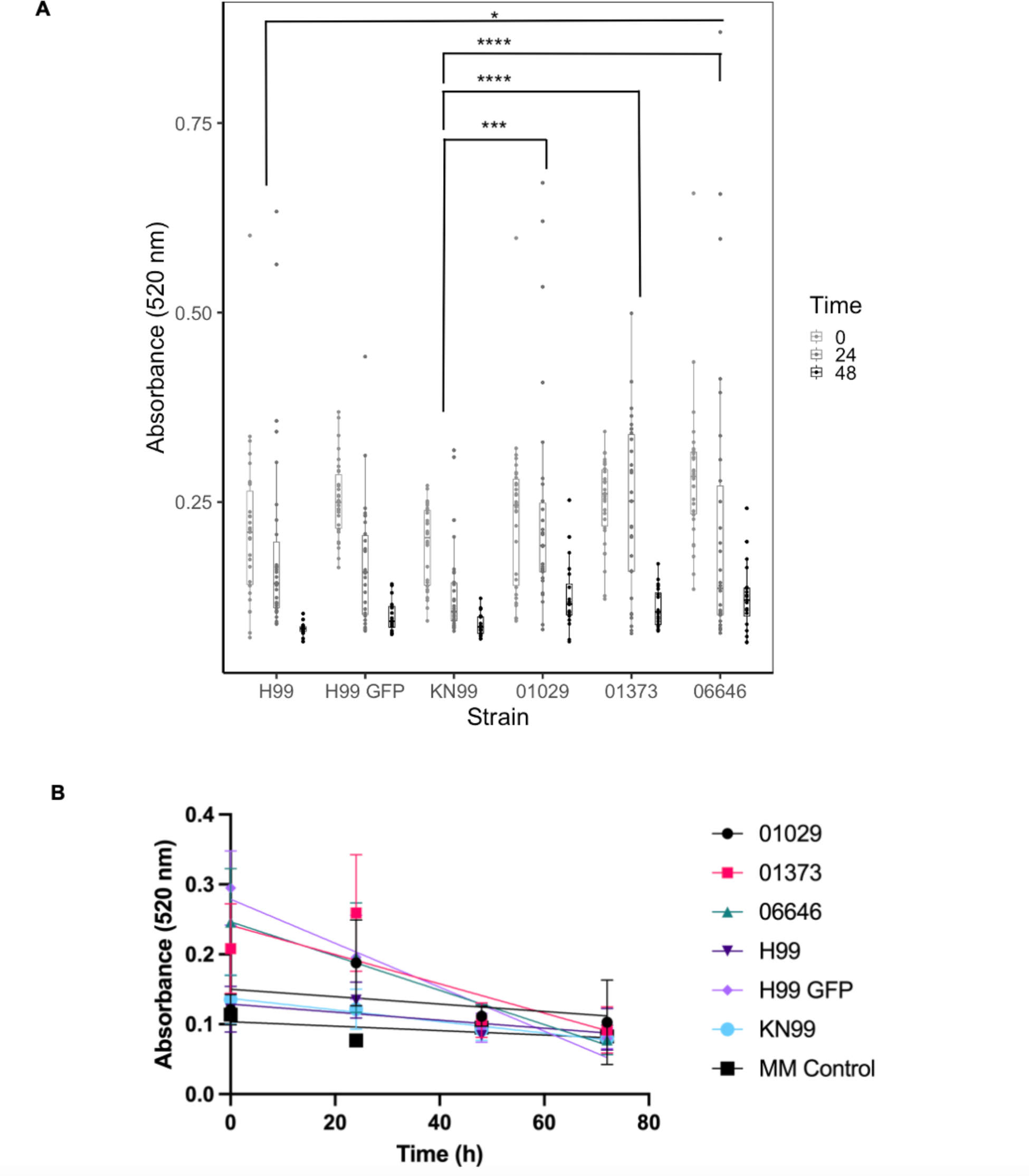
Optimized 24h colorimetric assay with comparisons between H99, H99 GFP, KN99α, and KN99α mutant strains. Complete color change from purple to light blue observed for every strain tested. **A.** Graphical results of comparison between different *C. neoformans* strains with 24 h assay. Each point represents an individual measurement. Overall significant differences between KN99α and 01029 (P = 0.0006), KN99α and 01373 (P = 0.00005), KN99α and 06646 (P = 0.000007), and H99 and 06646 (P = 0.02). Experiments conducted for four trials. *, **, ***, **** denote P < 0.05, 0.01, 0.001, 0.0001 respectively. **B.** Linear Regression Analysis of change in 520nm absorbance over time across *C. neoformans* strains with 24 h laccase assay method. Each point represents an average of measurements. As such, the following comparisons were statistically significant: H99 and H99 GFP (P << 0.00001), H99 and 01373 (P < 0.05), H99 and 06646 (P << 0.00001), H99 GFP and KN99α (P << 0.00001), H99 GFP and 01029 (P << 0.00001), KN99α and 06646 (P << 0.00001), 01029 and 06646 (P < 0.01). This statistical analysis was corrected for multiple comparisons.

**Figure 7.**
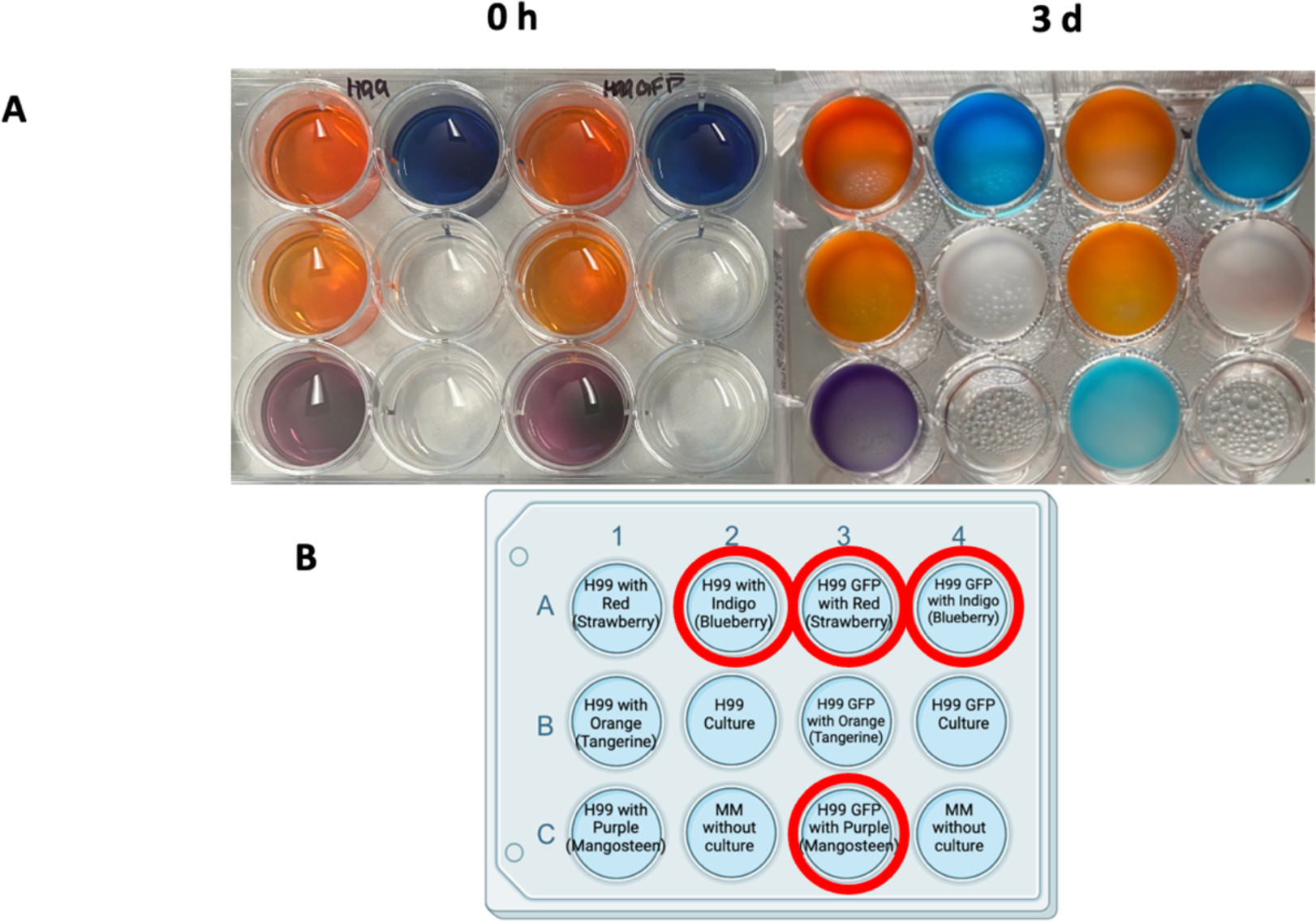
Laccase activity observed in H99 GFP strain after 3 d with various food dye colors. Color change observed with red color degradation: oranges to lighter orange hues, mangosteen (purple) to blue, and indigo to lighter blue. **A.** Left: 0 h photo of plate with H99 and H99 GFP wells with different colors. Right: 3 d photo of plate after being left on shaker in 30°C incubator. Experiment conducted in triplicate. **B.** Diagram of 12-well plate set up diagram for comparison between H99 and H99 GFP strain with the food coloring colors used, official names given by Limino brand in parentheses. Wells with a red circle designation are colors that showed the red color degradation.

**Figure 8.**
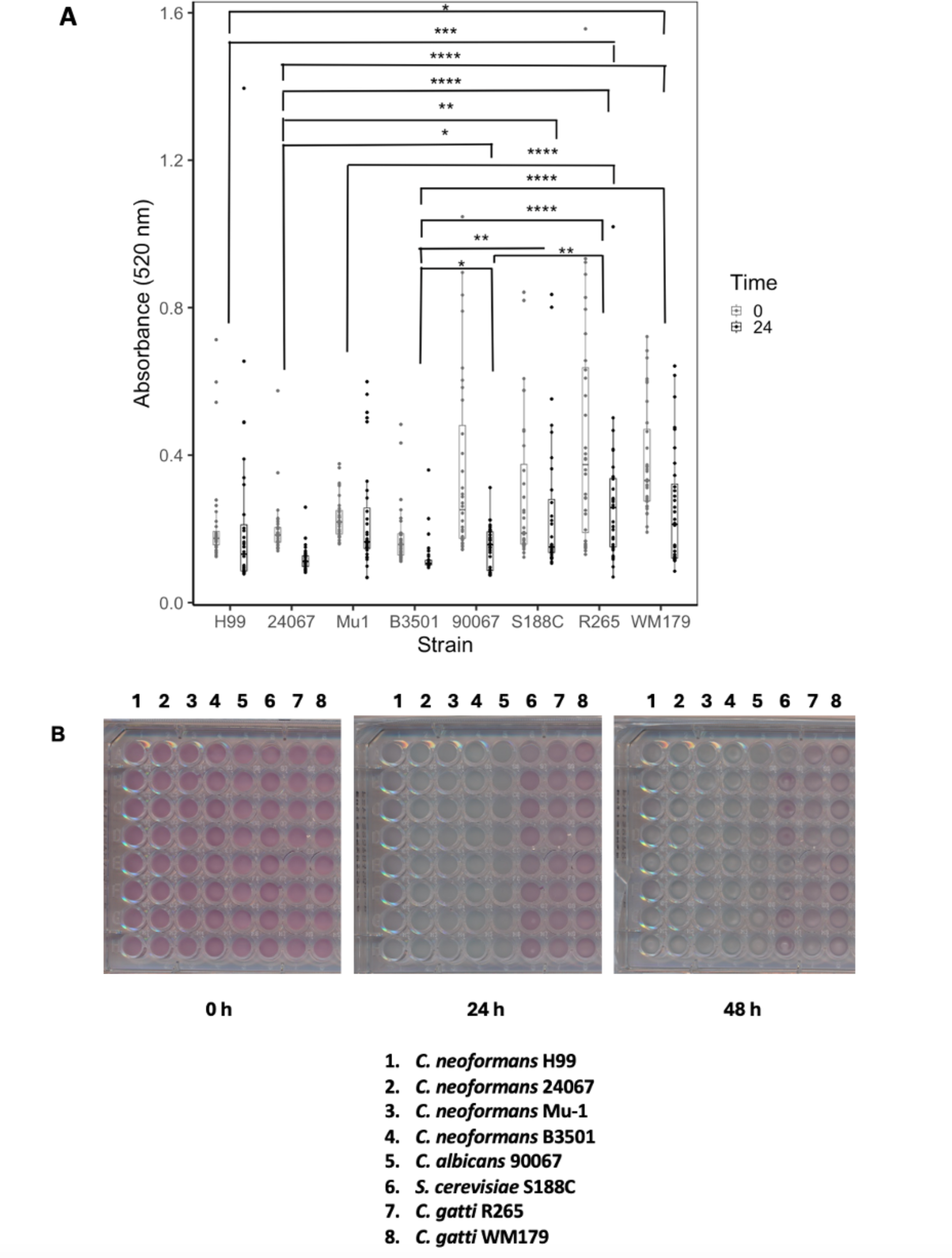
Optimized 24 h colorimetric assay with comparisons between *C. neoformans* and other fungal strains. **A.** Graphical results of comparison between different fungal strains with 24 h assay. Each point represents an individual measurement. Statistical analyses were conducted to compare the absorbance measurements across 24 h. Overall significant differences between H99 and *C. gatti* R265 (P = 0.00004), H99 and *C. gatti* WM179 (P = 0.02), 24067 and *C. albicans* 90067 (P = 0.04), 24067 and *C. gatti* R265 (P = 0.0000000), 24067 and *S. cerevisiae* S188C (P = 0.006), 24067 and *C. gatti* WM179 (0.000004), B3501 and *C. albicans* 90067 (P = 0.02), B3501 and *C. gatti* R265 (P = 0.0000000), B3501 and *S. cerevisiae* S188C (P = 0.002), B3501 and *C. gatti* WM179 (P = 0.000001), Mu-1 and *C. gatti* R265 (P = 0.0001), and *C. gatti* R265 and *C. albicans* 90067 (P = 0.005). Experiments conducted for four trials. *, **, ***, **** denote P < 0.05, 0.01, 0.001, 0.0001 respectively. **B**. Mangosteen (Purple) to blue color change observations. Images of plates comparing degradation of Mangosteen color over 48 h across fungal strains listed in the figure. Experiment conducted in triplicate.

With the combined data containing all individual trials of the first set of experiments comparing absorbance at 520 nm across different *C. neoformans* strains, we observed overall significant differences in absorbance between KN99α and 01029 (P = 0.0006), KN99α and 01373 (P = 0.00005), KN99α and 06646 (P = 0.000007), and H99 and 06646 (P = 0.02). In one of our trials, we observed significant differences in absorbance between the KN99α strain and its mutant strains 01029 (P = 0.009), 01373 (P = 0.003), and 06646 (P = 0.03). Significant differences between the strains tested within another trial are described in the figure legend of **Figure 6B**. While we observed a more noticeable difference in color change between the H99 and H99 GFP strain **(Figure 7)**, this was not always reflected in the statistical analyses particularly with the combination of multiple trials, possibly due to a loss of power with the analysis.

For the next group of experiments comparing absorbance at 520 nm across both *C. neoformans* strains and different yeast strains, we observed significant differences in absorbance measurements between H99 and *C. gatti* R265 (P = 0.00004), H99 and *C. gatti* WM179 (P = 0.02), 24067 and *C. albicans* 90067 (P = 0.04), 24067 and *C. gatti* R265 (P = 0.0000000), 24067 and *S. cerevisiae* S188C (P = 0.006), 24067 and *C. gatti* WM179 (0.000004), B3501 and *C. albicans* 90067 (P = 0.02), B3501 and *C. gatti* R265 (P = 0.0000000), B3501 and *S. cerevisiae* S188C (P = 0.002), B3501 and *C. gatti* WM179 (P = 0.000001), Mu-1 and *C. gatti* R265 (P = 0.0001), and *C. gatti* R265 and *C. albicans* 90067 (P = 0.005).

### Advantages of food dye based colorimetric assay relative to ABTS assay

We compared the standard ABTS assay to our colorimetric assay. The H99 GFP well with the higher concentration of culture showed a color change within 5 minutes of adding the ABTS solution to the well, so our initial photograph shows this color change already **(Figure 9A).** However, this color change was impermanent since it was not clearly evident after 24 h. This places a limitation on the ABTS experiment because it is possible to miss the window of color change and not obtain the absorbance measurements or photographs needed. Additionally, with our ABTS assay we observed the clearest color change mostly within the H99 GFP strain and not the H99 strain **(Figure 9A).** With the 24h colorimetric assay, we can observe a clear color change within 24 h for multiple *C. neoformans* strains and this color change exhibits permanence allowing for absorbance measurements or photographs to be taken post-color change at the investigator’s convenience. Additionally, there is a significant difference in cost of materials for the ABTS assay compared to the food coloring assay. ABTS solution is sold by Roche® Life Science Products in a quantity of 300 mL for $349.00 USD at the time of this study^16^. The food coloring used in this experiment is available on Amazon.com, costing $2.66/ Fl oz^17^. One fluid ounce is equivalent to 29.57 mL, so purchasing an equivalent amount of about 300 mL of food coloring would cost about $26.99, showing that the food coloring method is a significantly cheaper way to confirm laccase activity^18^.

**Figure 9.**
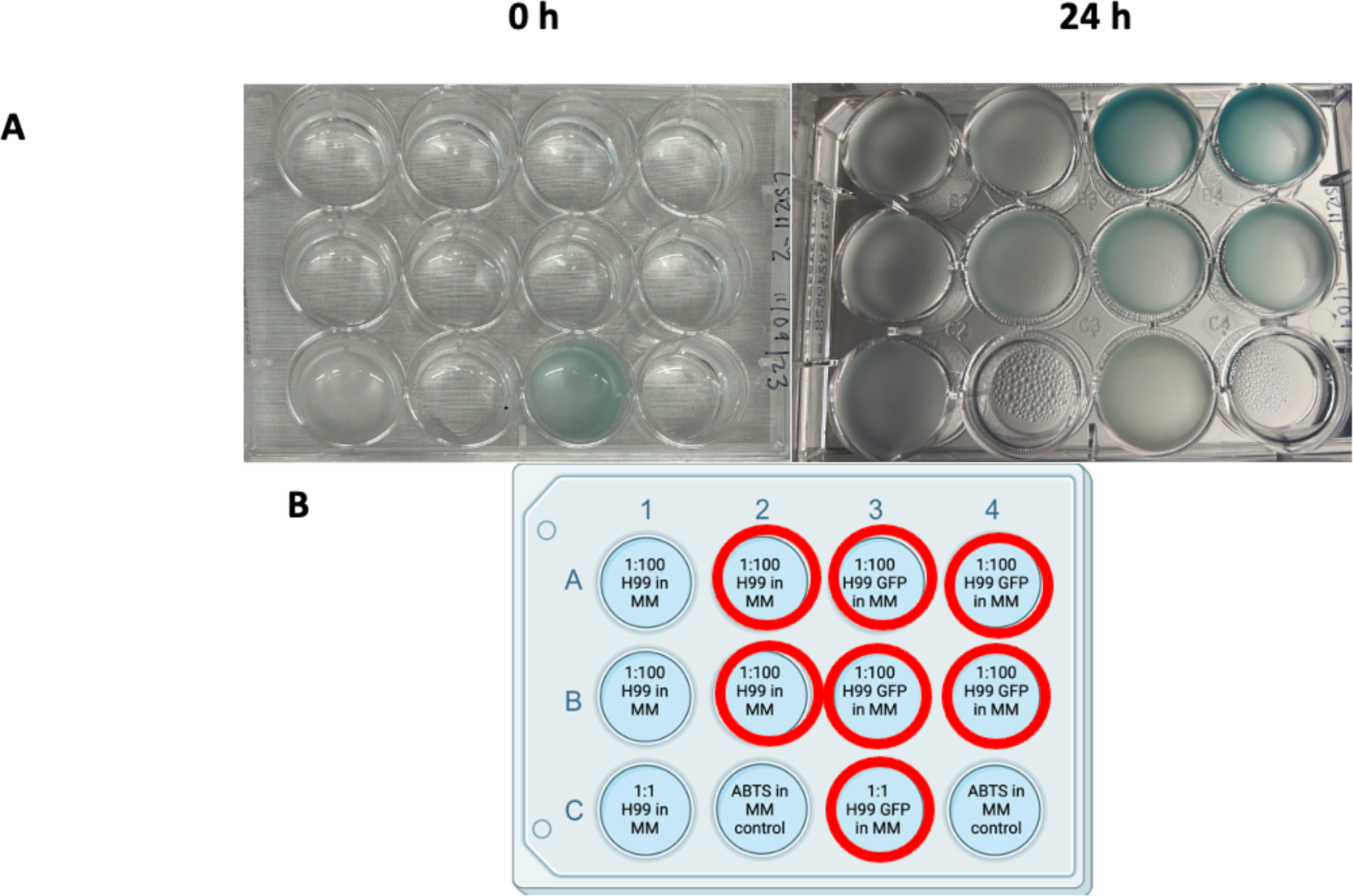
ABTS assay. Color change observed from clear to shades of blue-green. **A.** Left: Initial photo of ABTS assay plate at 0 h. Right: Photo of ABTS assay plate after 24 h in 30°C incubator on shaker. Note that well 3C turned blue rapidly but by 24 h the color was largely gone. **B.** 12-well plate set up diagram of ABTS experiment, created with Biorender.com. Wells with a red circle designation are where a color change from clear to shades of blue-green was observed.

## Discussion

Laccase enzymes are involved in the survival strategies of *C. neoformans* through the process of melanization. Melanization protects fungal cells from reactive oxidative stress, among other stressors, and investigation of the laccase enzyme role in *C. neoformans* pathogenesis can provide insight into the fungal survival strategy. Laccases are also relevant in many aspects of the food industry and may be useful in reducing the environmental impact of synthetic dyes from factory production. Synthetic food dyes may be used as an efficient and cheaper assay of laccase activity in *C. neoformans* cultures, and a method to quantify laccase activity is of interest in a variety of fields ranging from environmental concerns from accumulation of dyes in bodies of water, and the defense mechanisms of the pathogenic fungi^19, 20^. Since laccase is involved in catalyzing the melanization defense mechanism of *C. neoformans*, the relative comparisons of laccase activity across *C. neoformans* strains could be valuable in understanding differences across different strains.

Our results suggest that laccase irreversibly breaks down the red pigment known as Food Red 7 and Acid Red 27, as color changes were not observed with the *lac1*Δ mutant strain. We hypothesize this phenomenon is due to the enzyme’s ability to catalyze formation of free radicals through “removal of a hydrogen atom from the hydroxyl group of ortho- and para-substituted mono- and polyphenolic substrates”^19^. We initially considered that agitation was necessary to promote red color degradation in the wild-type H99 strain since the cryptococcal laccase reaction uses oxygen, but found that it was not necessary since plates at room temperature without agitation still showed color change when exposed to elevated glucose conditions albeit requiring a few more days to exhibit color degradation.

We observed more colorimetric activity in cultures with higher concentrations of glucose. This result was unexpected, as previous literature reported increased melanization at lower glucose concentrations which we expected to correlate with laccase activity^21^. In fact, a review of laccase activity cites glucose as a repressor^2^. However, those observations refer to melanization and not to direct laccase activity^21^. However, melanization may not always be a reliable proxy measurement for laccase activity^22^. Other reports have noted that laccase expression in *C. neoformans* is induced during glucose starvation, but also “stimulated by copper, iron, and calcium and repressed at elevated temperatures.”^23^. The glucose-dependency of color degradation shown in these experiments leads us to hypothesize that the phenomenon we observed could require energy. Alternatively, the glucose effect could reflect changes in the reduction potential of the solution that potentiate the dye destroying reaction of laccase. The laccase reaction is heavily influenced by the reduction potential of the solution and glucose has reducing properties as illustrated by the blue bottle experiment done in high school chemistry courses^24, 25^.

When comparing different *C. neoformans* strains, we found that the H99 GFP strain showed signs of increased laccase activity with an earlier visible color change and a greater change in absorbance at 520nm over time. We don’t have an explanation for this phenomenon but suspect that linking the GFP construct to the actin promoter of this *C. neoformans* laboratory strain could be causing secondary metabolic effects. Additionally, we note that GFP can generate free radicals and affect the oxidative state of the cell^26^. Given the importance of reduction potential in the laccase reaction, it is possible that the enhanced color associated with GFP expressing *C. neoformans* reflects altered oxidative conditions in the cells^24^.

The 24-hour color change method requires absorbance measurements to quantify differences in the extent of red color degradation by the fungi. Therefore, we have determined that the different variations of the assay could be used for different purposes depending on what is needed for the experiment. Larger wells and higher cell densities can be used to observe color change over the course of 3-7 days to provide a positive or negative conclusion on whether there is laccase activity, whereas resuspending smaller volume cultures in high glucose MM can determine reaction differences over the course of 24 h. Strain differences did not always reach statistical significance even though color changes were visible to the human eye, and this could reflect variability sensitivity of the plate reader used and a need to further optimize the assay to improve quantification of results. While significant differences were observed when comparing *C. neoformans* strains to other fungal strains, the relevance of these results in terms of laccase expression is unknown. As of now, this assay is preferentially suited for determining whether a strain produces laccase rather than comparing strain activities.

The absorbance spectrum revealed a new peak at ∼420 nm in wells containing *C. neoformans* that degraded the food coloring, suggesting the formation of a new product when pre-treated with antioxidant. This new product was chemically stable. Filamentous fungi have been found to degrade a red diazo dye, using laccases to transfer the azo dye to nontoxic products^27^. On the other hand, synthetic dyes that are made up of aromatic compounds, such as azo dyes, when degraded produce amines which are mutagenic to humans^28^. These studies suggest that these could be products that were produced in our experiments, but further investigation is needed to determine what product was produced by the fungal cells when degrading the food coloring.

Currently, the most widely used laccase activity screen in the cryptococcal field is the 2,2′-azino- bis (3-ethylbenzothiazoline-6-sulfonic acid) (ABTS) assay in which blue-colored ABTS is oxidized to green-colored ABTS+. A disadvantage of this assay is that the ABTS reduction is impermanent, and samples must be measured before the oxidized ABTS+ is reduced. In contrast, the observed color change in our assay was permanent with no time-sensitive window required for doing the measurement. Samples may be left for long periods of time without special preservation before measurement. We also observed differences in the degree of visible color change across strains in the ABTS assay, which so far have been mitigated by the 24-hour colorimetric assay that shows complete color change across the strains we tested. The colorimetric assay with food coloring materials is also significantly less costly relative to the cost of ABTS solution^16^.

The *C. neoformans* laccase enzyme has other functions in addition to making melanin. For example, *C. neoformans* laccase is involved in the production of fungal prostaglandins^4^. It is possible that laccase deactivates molecules toxic to the fungi, and that its effect on food dye color is a reflection of its non-specific chemical activity. In this study we have used this effect to develop a new assay for laccase activity that can be used to study fungal laccases.

## Acknowledgements

Figure 2D, Figure 7B, Figure 9B, and Supplemental Figure 1 were created with Biorender.com. A.C. was supported in part by NIH grants AI162381, AI152078, and HL059842. Funders played no role in the experimental design or outcome of the project.

---

**Supplemental Table 1:** Limino Food Coloring Ingredients.

| **Color Name** | **Food Coloring Components** | **CAS No.** |
| --- | --- | --- |
| Blueberry | Acid Red 27 | 915-67-3 |
|  | Food Blue No. 1 | 3844-45-9 |
| Lemon | Food Yellow No. 4 | 1934-21-0 |
|  | Food Yellow 3 | 2783-94-0 |
| Lime | Food Yellow No. 4 | 1934-21-0 |
|  | Food Blue No. 1 | 3844-45-9 |
| Mangosteen | Acid Red 27 | 915-67-3 |
|  | Food Blue No. 1 | 3844-45-9 |
| Purple Cabbage | Food Blue No. 1 | 3844-45-9 |
|  | Food Yellow 3 | 2783-94-0 |
|  | Acid Red 27 | 915-67-3 |
| Strawberry | Food Red 7 | 2611-82-7 |
|  | Food Yellow 3 | 2783-94-0 |
| Tangerine | Food Yellow 3 | 2783-94-0 |
|  | Food Red 7 | 2611-82-7 |

Ingredients added to the mixture: Sorbitol (CAS No. 50-70-4), Water (CAS No. 7732-18-5), Glycerin (CAS No. 56-81-5), Carboxymethylcellulose sodium (CAS No. 9004-32-4), Potassium sorbate (CAS No. 590-00-1)

**Supplemental Figure 1:**
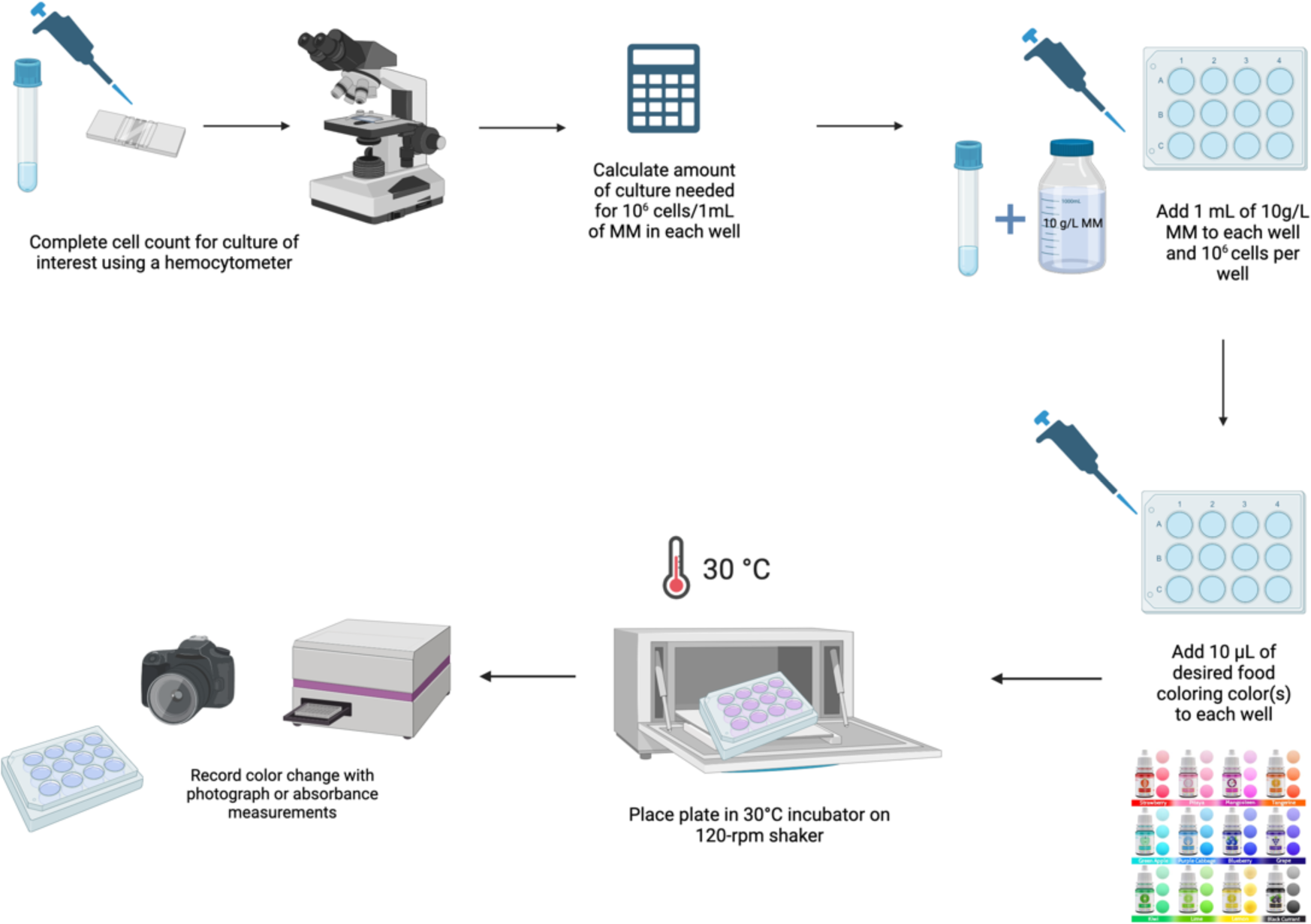
Lightweight Laccase Activity Assay. Lightweight Laccase Activity Assay, Image created with Biorender and photo of food coloring from Limino.

**Supplemental Figure 2:**
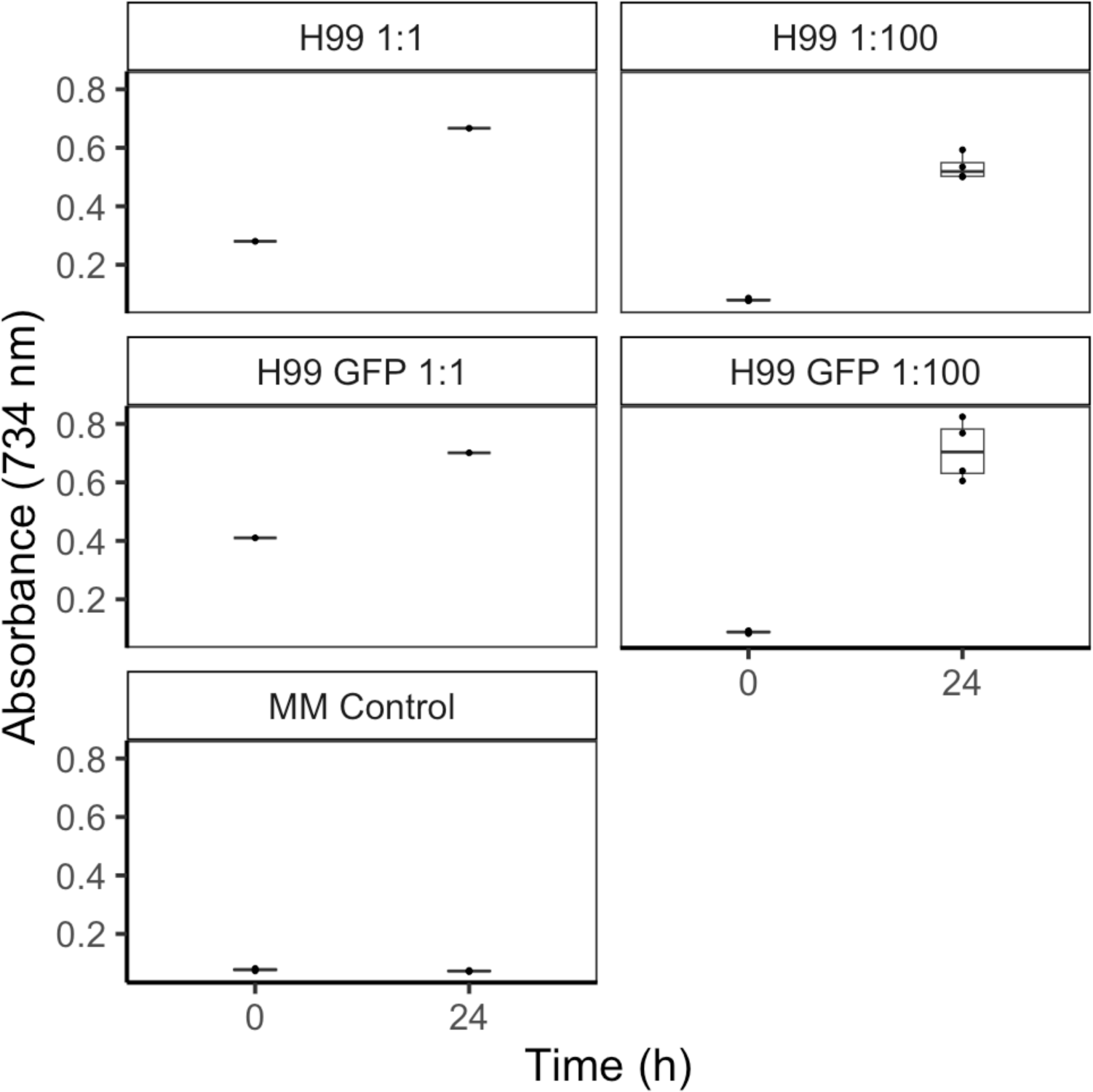
Difference in Absorbance over time with ABTS Assay. Difference in absorbance at 734nm wavelength for H99 and H99 GFP cultures in different dilutions in MM with ABTS. Each point represents an individual measurement.

